# Ascorbate depletion increases quiescence and self-renewal potential in hematopoietic stem cells and multipotent progenitors

**DOI:** 10.1101/2024.04.01.587574

**Authors:** Stefano Comazzetto, Daniel L. Cassidy, Andrew W. DeVilbiss, Elise C. Jeffery, Bethany R. Ottesen, Amanda R. Reyes, Sarah Muh, Thomas P. Mathews, Brandon Chen, Zhiyu Zhao, Sean J. Morrison

**Affiliations:** Children’s Research Institute and the Department of Pediatrics, University of Texas Southwestern Medical Center, Dallas, TX 75390, USA; Howard Hughes Medical Institute, University of Texas Southwestern Medical Center, Dallas, TX 75390, USA

**Keywords:** quiescence, metabolism, vitamin C, bone marrow transplantation

## Abstract

Ascorbate (vitamin C) limits hematopoietic stem cell (HSC) function and suppresses leukemia development by promoting the function of the Tet2 tumor suppressor. In humans, ascorbate is obtained from the diet while in mice it is synthesized in the liver. In this study, we show that deletion of the Slc23a2 ascorbate transporter severely depleted ascorbate from hematopoietic cells. *Slc23a2* deficiency increased HSC reconstituting potential and self-renewal potential upon transplantation into irradiated mice. *Slc23a2* deficiency also increased the reconstituting and self-renewal potential of multipotent hematopoietic progenitors (MPPs), conferring the ability to long-term reconstitute irradiated mice. *Slc23a2*-deficient HSCs and MPPs divided much less frequently than control HSCs and MPPs. Increased self-renewal and reconstituting potential were observed particularly in quiescent *Slc23a2*-deficient HSCs and MPPs. The effect of *Slc23a2* deficiency on MPP self-renewal was not mediated by reduced Tet2 function. Ascorbate thus regulates quiescence and restricts self-renewal potential in HSCs and MPPs such that ascorbate depletion confers MPPs with long-term self-renewal potential.

**KEY POINTS:**

- Deletion of the ascorbate transporter, Slc23a2, increases quiescence and self-renewal potential in HSCs and multipotent progenitors
- Ascorbate depletion is sufficient to confer long-term self-renewal potential upon multipotent hematopoietic progenitors

## INTRODUCTION

The daily production of blood cells in mammals is sustained by hematopoietic stem cells (HSCs), multipotent hematopoietic progenitors (MPPs), and restricted hematopoietic progenitors that reside in the bone marrow. HSCs have long-term self-renewal potential and give long-term multilineage reconstitution upon transplantation into irradiated mice^1–3^. MPPs persist and contribute to hematopoiesis over long periods of time in normal adult mice^4–8^ but when small numbers of MPPs are transplanted into irradiated mice they exhibit limited self-renewal potential and only transiently reconstitute^9–11^. HSCs appear to undergo a limited number of cell divisions before being fated to differentiate as quiescent HSCs have more reconstituting potential and more self-renewal potential than dividing HSCs^12,13^. Adult HSCs that have undergone multiple rounds of cell division have little reconstituting potential^14,15^.

Few mechanisms that limit the self-renewal potential of MPPs have been identified. Deficiency for Jumonji and AT-rich interaction domain containing 2 (Jarid2), a transcriptional repressor, confers long-term reconstituting potential upon MPPs, partly by increasing the expression of *Runx1t1* and *Mycn*^16^. Deficiency for *Trp53* and *Cdkn2a* increases the reconstituting potential of HSCs and confers long-term reconstituting potential upon MPPs^17^.

Metabolomic analysis showed that HSCs, MPPs, and restricted hematopoietic progenitors are metabolically distinct^18–21^. Ascorbate (vitamin C) is obtained from the diet in humans but synthesized in the liver of mice^22^. Ascorbate uptake by hematopoietic cells is mediated by the Slc23a2 transporter^18,23^, which is highly selective for ascorbate^24–27^. Ascorbate levels are high in HSCs and MPPs but decline with differentiation^18^. In a prior study, we systemically depleted ascorbate in *Gulonolactone oxidase* (*Gulo*) deficient mice, which lack the ability to synthesize ascorbate^18^. Systemic ascorbate depletion increases the reconstituting potential of HSCs and promotes the development of leukemia, largely by reducing the function of Tet2, a tumor suppressor that demethylates DNA^18^. Ascorbate is also necessary for T cell, plasma B cell, and erythrocyte differentiation^18,23,28–30^. Other essential dietary nutrients also regulate HSC function, including vitamin A^20,31^ and valine^32^. However, in contrast to ascorbate, dietary depletion of vitamin A or valine undermines HSC function.

In the current study we tested the effect of severe ascorbate depletion in hematopoietic cells by deleting the Slc23a2 transporter from hematopoietic cells. These mice do not develop scurvy because they have normal Slc23a2 function and ascorbate levels in non-hematopoietic cells. We found that ascorbate depletion in hematopoietic cells increased quiescence, reconstituting activity, and self-renewal potential in HSCs and MPPs. Ascorbate is thus not required for HSC or MPP function but promotes cell division under steady-state conditions and negatively regulates self-renewal potential.

## METHODS

### Mice

*Slc23a2* floxed mice (*Slc23a2^FL^)* were generated in a C57Bl/Ka background by introducing LoxP sites around exon 5 (which encodes part of the ascorbate transporter domain) using CRISPR-Cas9 (Supplemental Figure 1A-C). Cre-mediated recombination of this allele deletes exon 5, leading to a frameshift that eliminates Slc23a2-mediated ascorbate uptake. *Slc23a2^FL/^*^+^ mice were backcrossed at least 3 times onto a C57Bl/Ka background. *Mx1-Cre* mice^33^ were obtained from Jackson Laboratory (RRID:IMSR_JAX:003556). *Tet2^FL^* mice^34^ (RRID:IMSR_JAX:017573) were obtained from Dr. Iannis Aifantis and *Col1a1-H2B-GFP;Rosa26-M2-rtTA* mice^15^ (RRID:IMSR_JAX:016836) were obtained from Dr. Hanno Hock. All mice were maintained on a C57Bl/Ka background. We used 8-12 week-old littermates or age-matched mice in all experiments. To induce Cre recombinase expression, *Mx1-*Cre mice were given 3 intraperitoneal injections over 5 days of 40µg of polyinosinic-polycytidilic acid (poly I:C) (GE Healthcare) dissolved in PBS. Mice were housed in AAALAC-accredited, specific-pathogen-free animal care facilities at UT Southwestern Medical Center (UTSW). All procedures were approved by the UTSW Institutional Animal Care and Use Committee.

### Flow cytometric analysis and sorting of hematopoietic cells

Bone marrow was flushed from tibias and femurs using staining medium (Hanks Buffered Saline Solution (HBSS) supplemented with 2% bovine serum) and dissociated into a single cell suspension by gently triturating with a 23 gauge needle then filtering through a 40μm cell strainer. Cells were counted, and then stained with antibodies at 4°C for 30 minutes (except for anti-CD34 that was incubated for 90 minutes). For staining of HSCs and hematopoietic progenitors, cells were stained with antibodies against lineage markers (FITC- or APC-conjugated antibodies against CD2, CD3, CD5, CD8a, B220, Ter119, and Gr1), washed with staining medium and then resuspended in fluorophore-conjugated antibodies against c-kit, Sca1, CD150, CD48, CD16/32, Flt3, IL7Rα, CD105, and CD41. For analysis of lymphoid and myeloid cells, cells were stained with fluorophore-conjugated antibodies against Mac-1, Gr-1, B220 and CD3. Dead cells were identified and gated out of all analyses by staining with DAPI or with propidium iodide after antibody staining. Cells were analyzed using a FACSAria II (BD Biosciences), a FACSAria Fusion (BD Biosciences), or a FACS Lyric cytometer.

To sort HSCs, MPPs, HPC1 cells and HPC2 cells, tibias, femurs, pelvises, and vertebrae were crushed using a mortar and pestle. Cells were resuspended in staining medium and filtered through a 40μm cell strainer. The cells were stained with APC-efluo780 conjugated anti-c-kit antibody and c-kit^+^ cells were enriched using anti-APC paramagnetic microbeads (Miltenyi Biotec). Cells were stained with fluorophore-conjugated antibodies against lineage markers (CD2, CD3, CD5, CD8a, B220, Ter119, and Gr1), Sca1, CD150, and CD48. Cells were isolated by two successive rounds of sorting to ensure purity using a FACSAria II cytometer. The markers used for the identification and isolation of each cell population are listed in Supplemental Table 1 and representative flow cytometry gates are shown in supplemental Figure 1D-F.

### Bone marrow reconstitution assays

Recipient (CD45.1/CD45.2) mice were irradiated using an XRAD 320 X-ray irradiator (Precision X-Ray Inc.) with two doses of 540 rad at least 4h apart. For whole bone marrow transplantation, 500,000 unfractionated bone marrow cells from donor (CD45.2) mice and 2 million unfractionated bone marrow from competitor (CD45.1) mice were mixed and injected intravenously through the retro-orbital venous sinus. For HSC, MPP, HPC1 cell, and HPC2 cell transplants, the number of cells indicated in each experiment were mixed with 350,000 unfractionated bone marrow cells from competitor (CD45.1) mice and injected intravenously in the retro-orbital venous sinus. For secondary transplantation assays, 5 million unfractionated bone marrow cells from primary recipient mice were injected intravenously into irradiated secondary recipient mice. Every 4 weeks until at least 16-20 weeks after transplantation, blood was collected from the tail vein and subjected to ammonium-chloride potassium chloride red cell lysis. Cells were then stained with antibodies against CD45.1, CD45.2, Mac-1, Gr1, B220, and CD3 to evaluate the levels of donor cell engraftment in the myeloid, B, and T cell lineages. Recipient mice were considered long-term reconstituted if donor myeloid cells were present for at least 16 weeks after transplantation (>0.5% of myeloid cells were donor-derived).

### 5-bromo-2’-deoxyuridine incorporation

For analysis of the rates of HSC and MPP cell division, mice were intraperitoneally injected with 0.1mg/g body mass of 5-bromo-2’-deoxyuridine (BrdU), then administered water supplemented with 1mg/ml BrdU for 3 days before being killed to extract bone marrow for HSC and MPP isolation as described above. For analysis of the rates of HPC1, HPC2, and c-kit^+^ cell division, mice were intraperitoneally injected with 0.1mg/g body mass of BrdU and killed 2 hours after injection. Staining for BrdU incorporation into cells was performed using the BD APC BrdU Flow Kit following the manufacturer’s instructions. BrdU incorporation into each cell population was analyzed using a FACSAria II or a FACSAria Fusion (BD Biosciences) flow cytometer.

### Doxycycline administration for H2B-GFP pulse-chase experiments

*Col1a1-H2B-GFP;Rosa26-M2-rtTA;Slc23a2^FL/FL^;Mx1-Cre* mice and *Col1a1-H2B-GFP;Rosa26-M2-rtTA;Slc23a2^FL/FL^* littermate controls were given 3 intraperitoneal injections of 40µg of poly I:C (GE Healthcare) dissolved in PBS over 5 days to induce Cre expression. One week after the last poly I:C injection, mice were given water with 0.2% doxycycline (Research Product International) and 1% Sucrose (Sigma-Aldrich) for a total of 6 weeks to induce H2B-GFP expression. Twelve weeks after completing doxycycline treatment, mice were killed and bone marrow was obtained for the isolation of HSCs and MPPs as described above, then H2B-GFP levels in the cells was quantitated by flow cytometry.

### Ascorbate measurement by LC-MS/MS

Two million whole bone marrow cells from *Slc23a2^FL/FL^;Mx1-Cre* and *Slc23a2^FL/FL^* control mice were washed twice with ice-cold PBS. Cells were then resuspended in 200ul of 5% metaphosphoric acid (Sigma-Aldrich) with 1mM EDTA (Sigma-Aldrich) and 10µM ^13^C_6_-ascorbate (Toronto Research Chemicals) then centrifuged for 30 minutes at 4°C. Blood was collected from the tail vein and centrifuged at 3000 x g for 10 minutes at 4°C. Plasma was then collected and mixed 1:1 with 10% metaphoshoric acid with 2mM EDTA and 20µM ^13^C_6_-ascorbate. This mixture was then centrifuged for 30 minutes at 4°C. The supernatant from bone marrow cells or plasma was then transferred to a new Eppendorf tube and store at -80°C until analysis. Ascorbate levels were quantified by liquid chromatography-tandem mass spectrometry (LC-MS/MS) using a SCIEX 6500+ QTrap mass spectrometer coupled to an Exion LC system. Ascorbate was chromatographically resolved using a Waters HSS T3 column (50mm x 2.1mm, 1.7um bead size) with a binary solvent system of 0.1% formic acid in water (solvent A) and 0.1% formic acid in acetonitrile (solvent B) flowed at 0.25ml/minute. Identification of ascorbate by mass spectrometry was achieved by operating the mass spectrometer in negative ionization mode with multiple reaction monitoring (MRM). Ascorbate was quantified using a transition of m/z 175 to m/z 115 (declustering potential (DP): -25V, collision energy (CE): -18V, collision exit potential (CXP): -10V). Ascorbate was structurally identified with a second transition of m/z 175 to m/z 87 (DP: -25V, CE: -28V, CXP: -10V). The spiked-in internal standard of ^13^C_6_-ascorbate was monitored with a single transition of m/z 181 to m/z 119 (DP: -25V, CE: -18V, CXP: -10V). All dwell times were set to 50 ms. The concentration of ascorbate was determined using MultiQuant (SCIEX) and a standard curve of commercially available ascorbate.

### RNA-Seq library preparation and data analysis

Two thousand to eight thousand HSCs or MPPs were sorted into tubes containing RLT buffer from the RNeasy Micro Kit (Qiagen). Total RNA was extracted according to the manufacturer’s instructions. RNA integrity and concentration were measured using a Bioanalyzer (Agilent). One to six nanograms of RNA were used to generate a cDNA library using the SMARTer Stranded Total RNA-Seq Kit v2 - Pico Input Mammalian (Takara). Samples were sequenced with a NextSeq 500 (Illumina) using 75bp single-end reads. The quality of RNA-seq raw reads was checked using FastQC 0.11.8. Raw reads were mapped to the Ensembl GRCm38 mouse reference genome using Salmon 1.4.0. Mapped reads were quantified using HTSeq 0.9.1. Quantified mapped reads were normalized and gene expression levels were measured as fragments per thousand exonic bases per million mapped reads (FPKMs) using DESeq2 1.30.0 with R 4.0.2. Differential expression was assessed using DESeq2 1.30.0. Gene ontology analysis was performed using ShinyGO 0.80.

### Statistical analysis

Data in all figures were obtained in at least two independent experiments using different mice, as indicated in each figure legend. Data are shown as mean±standard deviation. Flow cytometry data were analyzed using Flowjo v.10.9 (Flowjo). Before analyzing the statistical significance of differences between treatments, we tested for normality in data distributions using the D’Agostino-Pearson omnibus test for samples with n>8 or the Shapiro-Wilk normality test for smaller samples. We also tested for similar variability among treatments using an F-test (when comparing two samples) or Levene’s median tests (when comparing more than two samples). When data were normally distributed and had similar variability among treatments, we used parametric tests to assess statistical significance. When the data significantly deviated from normality or variability significantly differed among treatments, we log2-transformed the data. If the log2 transformed data were normally distributed and had similar variability among treatments, we performed parametric tests to assess the statistical significance of differences among treatments using the log2 transformed data. If the log2 transformed data were not normally distributed or did not have similar variability among treatments, we performed non-parametric tests on the non-transformed data.

When assessing the statistical significance of differences between two treatments, statistical analyses were performed using student t-tests, Mann-Whitney non-parametric tests, multiple student t-tests, or Mann-Whitney tests with Holm-Sidak corrections for multiple comparisons (Graphpad Prism v.10). When assessing the statistical significance of differences between more than two treatments, statistical analyses were performed using one-way ANOVAs with Tukey’s corrections for multiple comparisons or Kruskal-Wallis tests with Dunn’s corrections (Graphpad Prism v.10). For transplantation experiments, we used a non-parametric mixed effects model, nparLD, to test for differences in donor cell reconstitution by cells of different genotypes. All statistical tests we used were two-sided.

## RESULTS

### Ascorbate depletion increases reconstituting potential in transplantation assays

Since germline *Slc23a2* deficiency is embryonically lethal^35^, we created a floxed allele of *Slc23a2*, enabling conditional deletion from hematopoietic cells using Mx1-Cre (supplemental Figure 1A-C). Eight-week-old *Slc23a2^FL/FL^;Mx1-Cre* and *Slc23a2^FL/FL^* control mice were injected with polyinosinic-polycytidilic acid (poly I:C) to induce Cre expression and then analyzed two to four weeks later. *Slc23a2^FL/FL^;Mx1-Cre* mice exhibited a 12-fold depletion of intracellular ascorbate in bone marrow hematopoietic cells (Figure 1A) but normal ascorbate levels in blood plasma (Figure 1B). *Slc23a2* deficiency in hematopoietic cells did not lead to scurvy, which kills *Gulo*-deficient mice within six weeks in the absence of dietary supplementation^36^. The *Slc23a2*-deficient mice did not die or appear ill more than 4 months after Cre induction (Figure 1C).

**Figure 1:**
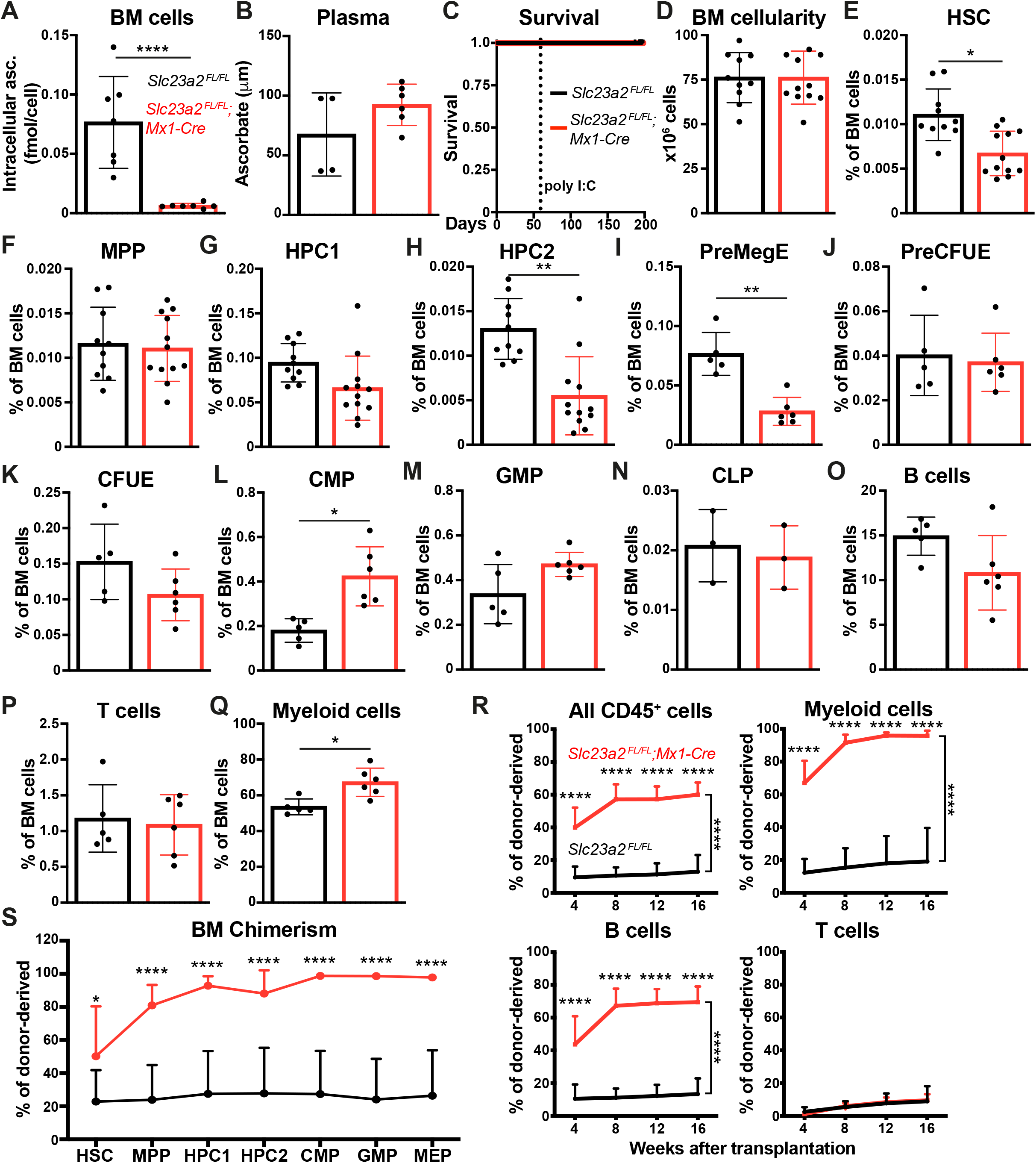
Ascorbate depletion increases the reconstituting potential of bone marrow cells upon transplantation into irradiated mice. (A-B) Ascorbate levels in bone marrow cells (A) and blood plasma (B) measured by mass spectrometry (a total of 4-7 mice per genotype in 2 independent experiments). (C) Kaplan-Meyer survival curve of *Slc23a2*-deficient and littermate control mice (a total of 8 mice per genotype in 3 independent experiments). (D-Q) Bone marrow cellularity from two tibias and two femurs (D) as well as the frequencies of HSCs (E), MPPs (F), restricted hematopoietic progenitors (G-N), and differentiated B (O), T (P), and myeloid (Q) cells in the bone marrow (a total of 3-12 mice per genotype from 6 independent experiments). (R-S) Donor cell reconstitution in the blood (R) and bone marrow (S) after transplantation of 500,000 donor bone marrow cells from *Slc23a2*-deficient or littermate control mice along with 2,000,000 competing wild-type cells into irradiated recipients (a total of 13-14 recipient mice per genotype transplanted with cells from 3 donors per genotype in 3 independent experiments). All data represent mean ± standard deviation. Each dot in panels A-Q represents a different mouse. Statistical significance was assessed using Student’s *t*-tests (A and D), Mann-Whitney tests (B and time points in R), Log-rank test (C), Student’s t-tests with Holm-Sidak’s correction for multiple comparisons (E-Q), nparLD non-parametric mixed models with Holm-Sidak’s multiple comparisons corrections to test differences in overall reconstitution (R), or Mann-Whitney tests with Holm-Sidak’s correction for multiple comparisons (S). All statistical tests were two-sided (* *P*<0.05; ** *P*<0.01; *** *P*<0.001; **** *P*<0.0001).

The flow cytometry gates and cell surface markers used to identify each cell population characterized in this study are shown in supplemental Figure 1D-F and supplemental Table 1, respectively. *Slc23a2* deficiency did not affect bone marrow cellularity (Figure 1D) or the frequencies of MPPs or HPC1 cells (Figure 1F-G), but it reduced the frequencies of HSCs (Figure 1E) and HPC2 cells (Figure 1H). *Slc23a2* deficiency also reduced the frequency of megakaryocytic-erythroid progenitors (PreMegEs) (Figure 1I), but it did not change the frequency of erythroid progenitors (PreCFUEs and CFUEs) (Figure 1J-K). *Slc23a2* deficiency increased the frequency of common myeloid progenitors (CMPs) (Figure 1L) but did not significantly affect the frequencies of granulocyte-monocyte progenitors (GMPs) or common lymphoid progenitors (CLPs) (Figure 1M-N). *Slc23a2* deficiency did not affect the frequencies of differentiated B or T cells in the bone marrow (Figure 1O-P), but it did increase the frequency of myeloid cells (Figure 1Q). Overall, ascorbate depletion in hematopoietic cells modestly depleted HSCs and increased myelopoiesis in the bone marrow.

To test if *Slc23a2* deficiency affects the reconstituting potential of bone marrow cells, we competitively transplanted whole bone marrow cells from *Slc23a2^FL/FL^;Mx1-Cre* and *Slc23a2^FL/FL^*control mice (2-4 weeks after poly I:C treatment) into irradiated recipient mice. Consistent with our prior study^18^, *Slc23a2*-deficient bone marrow cells gave significantly higher levels of donor myeloid and B cell reconstitution as compared to control cells (Figure 1R). *Slc23a2* deficient and control cells did not differ in terms of T cell reconstitution in these experiments, though this was not unexpected as ascorbate deficiency impairs T cell differentiation^23^. At 16 weeks after transplantation, we also examined donor cell chimerism among hematopoietic stem and progenitor cells in the bone marrow. *Slc23a2* deficient cells gave significantly higher levels of donor cell reconstitution of HSCs and all hematopoietic progenitor cell populations we examined as compared to control cells (Figure 1S). Therefore, ascorbate depletion increased the reconstituting potential of bone marrow cells in irradiated mice despite a reduction in HSC frequency in the bone marrow.

### Ascorbate depletion increased HSC and MPP self-renewal potential

To better understand the increased reconstituting potential of ascorbate depleted bone marrow cells, we competitively transplanted flow cytometrically isolated HSCs, MPPs, HPC1 or HPC2 cells from *Slc23a2^FL/FL^;Mx1-Cre* and *Slc23a2^FL/FL^* control mice (2-4 weeks after poly I:C treatment) into irradiated recipient mice. We first transplanted 25 donor HSCs per recipient from *Slc23a2^FL/FL^;Mx1-Cre* or littermate control mice along with 350,000 competitor bone marrow cells into irradiated recipient mice. Nearly all of the recipient mice were long-term multilineage reconstituted by donor cells: 14 of 17 recipients of control HSCs and 16 of 17 recipients of *Slc23a2*-deficient HSCs. As compared to control HSCs, *Slc23a2*-deficient HSCs gave significantly higher levels of donor myeloid and B cell reconstitution in the blood (Figure 2A) as well as donor HPC1 cell, CMP, GMP, and MEP reconstitution in the bone marrow (Figure 2B-C). We also serially transplanted whole bone marrow cells from these primary recipient mice into irradiated secondary recipients. The *Slc23a2*-deficient cells again gave significantly higher levels of donor myeloid and B cell reconstitution in the blood (Figure 2D). Six of 8 secondary recipients of control cells and 7 of 7 secondary recipients of *Slc23a2*-deficient cells were long-term multilineage reconstituted by donor cells. *Slc23a2*-deficient HSCs thus had increased reconstituting potential in both primary and secondary recipient mice.

**Figure 2:**
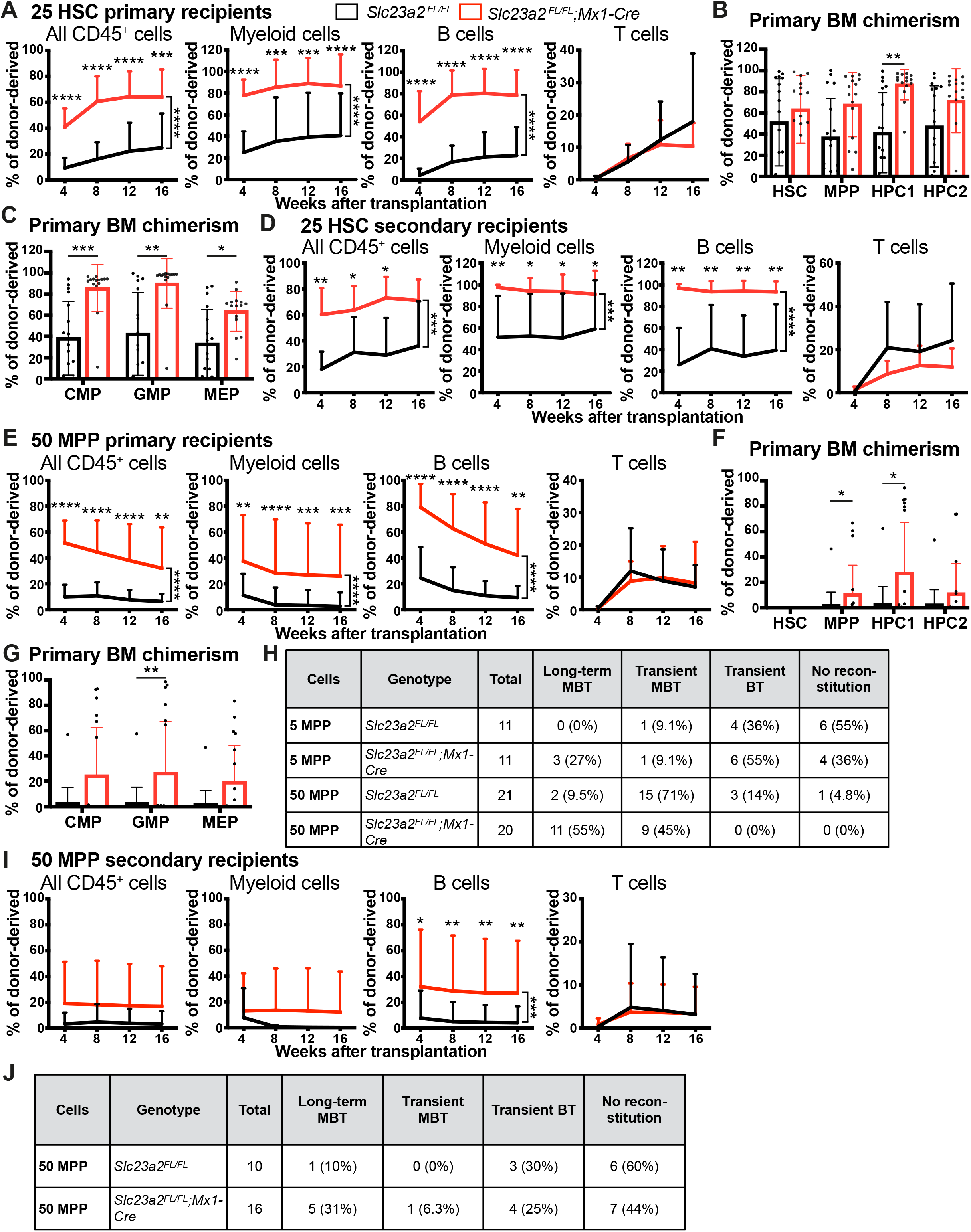
Ascorbate depletion increases the self-renewal potential of HSCs and MPPs. (A-C) Donor cell reconstitution in the blood (A) and bone marrow (B-C) of irradiated mice competitively transplanted with 25 HSCs from *Slc23a2*-deficient or littermate control mice (a total of 15-17 recipient mice per genotype transplanted with cells from 4 donors per genotype in 4 independent experiments). (D) Donor cell reconstitution in the blood of secondary recipients of 5 million bone marrow cells from the primary recipients in panel A (a total of 7-8 recipient mice per genotype transplanted with cells from 2 donor mice per genotype in 2 independent experiments). (E-G) Donor cell reconstitution in the blood (E) and bone marrow (F-G) of irradiated mice competitively transplanted with 50 MPPs isolated from *Slc23a2*-deficient or littermate control mice (a total of 20-21 recipient mice per genotype transplanted with cells from 5 donors per genotype in 5 independent experiments). (H) Summary of donor cell reconstitution profiles from primary recipients of MPPs. (I, J) Donor cell reconstitution in the blood of secondary recipients of bone marrow cells from the primary recipients in panel E (a total of 10-16 recipient mice per genotype transplanted with cells from 5 donor mice per genotype in 5 independent experiments). All data represent mean ± standard deviation. Each dot in panels B, C, F, and G represents a different mouse. Statistical significance was assessed using Mann-Whitney tests with Holm-Sidak’s corrections for multiple comparisons (B-C, F-G), Mann-Whitney tests to assess differences among genotypes at each timepoint and nparLD models with Holm-Sidak’s multiple comparisons corrections to test differences among overall reconstitution (A, D-E, I). All statistical tests were two-sided (* *P*<0.05; ** *P*<0.01; *** *P*<0.001; **** *P*<0.0001).

We next tested if *Slc23a2* deficiency altered the reconstituting potential of MPPs by competitively transplanting 50 MPPs from *Slc23a2^FL/FL^;Mx1-Cre* or *Slc23a2^FL/FL^* control mice into irradiated recipients. As compared to control MPPs, *Slc23a2*-deficient MPPs gave significantly higher levels of donor myeloid and B cell reconstitution in the blood (Figure 2E) as well as donor MPP, HPC1 cell, and GMP reconstitution in the bone marrow (Figure 2F-G). While most recipients of control MPPs were transiently reconstituted by donor cells (18 of 21 mice) most recipients of *Slc23a2*-deficient MPPs were long-term multilineage reconstituted by donor cells (11 of 20 mice) (Figure 2H). This suggested that ascorbate depletion conferred long-term self-renewal potential upon MPPs, which normally only give transient multilineage reconstitution^9,37^.

We also competitively transplanted 5 MPPs from *Slc23a2^FL/FL^;Mx1-Cre* or *Slc23a2^FL/FL^* mice into irradiated recipients. Three of 11 primary recipients of *Slc23a2*-deficient cells were long-term multilineage reconstituted by donor cells, while none of 11 primary recipients of control cells were long-term multilineage reconstituted by donor cells (Figure 2H). Limiting dilution analysis^38^ of the 5 and 50 cell transplant data indicated that *Slc23a2* deficiency increased the frequency of long-term multilineage reconstituting cells in the MPP population by 10 fold: 1 in 530 control MPPs and 1 in 52 *Slc23a2*-deficient MPPs gave long-term multilineage reconstitution (*P* = 1.1 x10^-4^). Ascorbate depletion thus conferred long-term self-renewal potential upon MPPs.

To further test this we serially transplanted bone marrow cells from recipients of 50 MPPs into irradiated secondary recipients. The *Slc23a2*-deficient cells again gave significantly higher levels of donor B cell reconstitution in the blood of secondary recipients as compared to control cells (Figure 2I). Only 1 of 10 (10%) secondary recipients of control cells were long-term multilineage reconstituted while 5 of 16 (31%) secondary recipients of *Slc23a2*-deficient cells were long-term multilineage reconstituted (Figure 2J).

Finally, we tested if *Slc23a2* deficiency altered the reconstituting potential of HPC1 or HPC2 cells by competitively transplanting 500 HPC1 cells or 200 HPC2 cells from *Slc23a2^FL/FL^;Mx1-Cre* or *Slc23a2^FL/FL^*control mice into irradiated recipients. These cell populations normally only transiently reconstitute irradiated mice^37,39^. *Slc23a2*-deficient HPC1 cells gave significantly higher levels of donor B cell reconstitution than control HPC1 cells (supplemental Figure 2A), though both gave only transient reconstitution (supplemental Figure 2B). *Slc23a2*-deficient HPC2 cells tended to give higher levels of donor cell reconstitution than control HPC2 cells, though the differences were mostly not statistically significant (supplemental Figure 2C). Again, *Slc23a2*-deficient and control HPC2 cells gave only transient reconstitution (supplemental Figure 2D). Thus, ascorbate depletion tended to increase the reconstituting potential of HPC1 and HPC2 cells but did not confer long-term self-renewal potential upon these progenitor populations.

### Ascorbate depletion reduced cell division by HSCs and MPPs

Although the markers we used to isolate HSCs and MPPs yield highly purified populations, these cells remain heterogeneous, particularly in terms of cell cycle regulation. To test if ascorbate depletion altered the composition of the HSC and MPP pools, we assessed CD229 and CD244, markers that distinguish functionally distinct subsets of HSCs and MPPs^37^. *Slc23a2* deficiency increased the frequency of HSC-1 (CD229^-^CD244^-^) cells while reducing the frequency of HSC-2 (CD229^+^CD244^-^) cells (Figure 3A-B). Since HSC-1 cells have increased self-renewal and reconstituting potential as compared to HSC-2 cells^37^, this reflected a shift toward more primitive HSCs in ascorbate depleted bone marrow. *Slc23a2* deficiency also increased the frequency of MPP-1 (CD229^-^CD244^-^) cells while reducing the frequencies of MPP-2 (CD229^+^CD244^-^) and MPP-3 (CD229^+^ CD244^+^) cells (Figure 3A-B). Since MPP-1 cells have increased reconstituting potential as compared to MPP-2 and MPP-3 cells^37^, this again reflected a shift toward more primitive MPPs in ascorbate depleted bone marrow.

**Figure 3:**
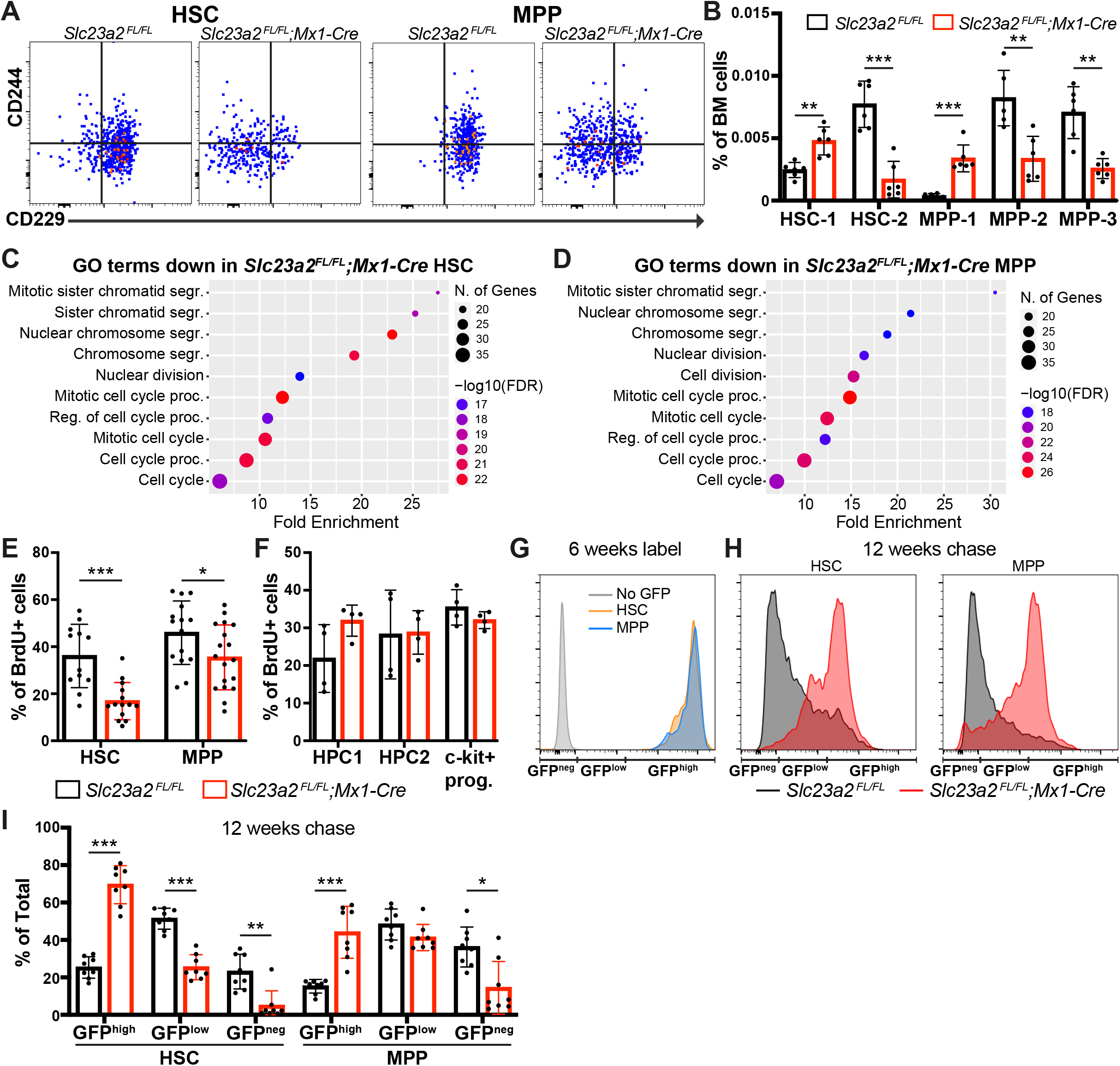
Ascorbate depletion promotes HSC and MPP quiescence. (A) Representative flow cytometry plots showing staining for CD229 and CD244 in HSCs and MPPs from *Slc23a2*-deficient or littermate control mice. These markers distinguish functionally distinct subsets of HSCs and MPPs that differ in terms of primitiveness^37^. (B) Frequencies of HSC-1, HSC-2, MPP-1, MPP-2, and MPP-3 subsets based on these markers (a total of 6 mice per genotype from 4 independent experiments). (C-D) Analysis of Gene Ontology (GO) biological process terms that were enriched among genes downregulated (FC<0.5 and FDR<0.01) in *Slc23a2*-deficient as compared to control HSCs (C) or MPPs (D) by RNA sequencing. (E-F) Incorporation of a 3 day (E) or 2 hour (F) pulse of BrdU into HSCs, MPPs, HPC1 cells, HPC2 cells, and all c-kit^+^ progenitors (a total of 4-18 mice per genotype from 3 (F) and 7 (E) independent experiments). (G) Representative H2B-GFP fluorescence in HSCs and MPPs from *Slc23a2^FL/FL^;Col1a1-H2B-GFP;Rosa26-M2-rtTA* mice without doxycycline treatment (grey) or after 6 weeks of doxycycline treatment (yellow HSCs, blue MPPs). (H) Representative H2B-GFP fluorescence in HSCs and MPPs from *Slc23a2*-deficient or littermate control mice after 12 weeks of chase without doxycycline. (I) The percentages of H2B-GFP^high^, H2B-GFP^low^, or H2B-GFP^neg^ HSCs and MPPs after 12 weeks of chase without doxycycline (a total of 8 mice per genotype analyzed in 4 independent experiments). All data represent mean ± standard deviation. Each dot represents a different mouse in panels B, E, F, and I. Statistical significance was assessed using Student’s *t*-tests with Holm-Sidak’s correction for multiple comparisons (B, E-F) or Mann-Whitney tests with Holm-Sidak’s corrections for multiple comparisons (I). All statistical tests were two-sided (* *P*<0.05; ** *P*<0.01; *** *P*<0.001).

To further explore this, we sequenced RNA from *Slc23a2*-deficient and control HSCs and MPPs. We identified 81 and 65 genes whose expression was significantly reduced as well as 9 and 19 genes whose expression was significantly increased in *Slc23a2*-deficient as compared to control HSCs and MPPs, respectively (Fold change >2 fold, FDR <0.01) (supplemental Table 2). Gene ontology (GO) term analysis revealed that all 10 of the most significantly downregulated GO terms in *Slc23a2*-deficient as compared to control HSCs and MPPs were related to cell division (Figure 3C-D). No GO terms were significantly enriched among the genes that were upregulated in *Slc23a2*-deficient as compared to control HSCs. This suggested that the main difference among *Slc23a2*-deficient as compared to control HSCs and MPPs was reduced cell division among the *Slc23a2*-deficient cells.

To test this directly we administered a 3 day pulse of 5-bromo-3-deoxyuridine (BrdU) to *Slc23a2^FL/FL^;Mx1-Cre* and *Slc23a2^FL/FL^*control mice then assessed BrdU incorporation into HSCs and MPPs by flow cytometry. Significantly lower percentages of *Slc23a2*-deficient HSCs and MPPs incorporated BrdU as compared to control HSCs and MPPs (Figure 3E). We also administered a 2 hour pulse of BrdU to *Slc23a2^FL/FL^;Mx1-Cre* and *Slc23a2^FL/FL^* control mice then assessed BrdU incorporation by HPC1 cells, HPC2 cells, and c-kit^+^ cells. We observed no significant difference in BrdU incorporation into *Slc23a2*-deficient as compared to control HPC1 cells, HPC2 cells, or all c-kit^+^ cells (Figure 3F). Ascorbate depletion thus selectively reduced the rate of cell division among HSCs and MPPs.

### Ascorbate depletion conferred long-term self-renewal potential upon quiescent MPPs

To more deeply assess the effect of ascorbate depletion on HSC and MPP cell division, we mated *Slc23a2^FL/FL^;Mx1-Cre* mice with *Col1a1-H2B-GFP;Rosa26-M2-rtTA* mice^15^ and then administered 6-weeks of doxycycline to label cells with H2B-GFP. We then withdrew the doxycycline so that the H2B-GFP levels would be diluted with each round of cell division over a subsequent 12 week chase (Figure 3G-H)^14,15^. After 12 weeks of chase, we quantified the percentages of HSCs and MPPs that had undergone few divisions (H2B-GFP^high^), an intermediate number of divisions (H2B-GFP^low^), or many divisions (H2B-GFP^neg^) (Figure 3G-H).

*Slc23a2* deficiency substantially increased H2B-GFP retention by both HSCs and MPPs, indicating much less cell division by these cells during the chase period. This was reflected by significantly higher percentages of H2B-GFP^high^ HSCs and MPPs and significantly lower percentages of H2B-GFP^low^ and H2B-GFP^neg^ HSCs and MPPs in *Slc23a2*-deficient as compared to control mice (Figure 3I). HSCs and MPPs thus undergo prolonged periods of quiescence after ascorbate depletion.

To test if reduced cell division was associated with increased reconstituting potential in HSCs, we competitively transplanted 20 H2B-GFP^high^ HSCs or 50 H2B-GFP^neg^ HSCs from *Slc23a2*-deficient and control mice after 12 weeks of chase into irradiated recipients. As expected^14,15^, among control cells, the H2B-GFP^high^ HSCs gave significantly higher levels of donor myeloid cell reconstitution and a higher percentage of recipients that were long-term multilineage reconstituted as compared to H2B-GFP^neg^ HSCs (Figure 4A-D). *Slc23a2*-deficient H2B-GFP^high^ HSCs gave significantly higher levels of donor myeloid and B cell reconstitution and a higher percentage of recipients that were long-term multilineage reconstituted as compared to control H2B-GFP^hi^ HSCs (Figure 4A-B). *Slc23a2*-deficient H2B-GFP^neg^ HSCs also gave significantly higher levels of donor myeloid, B, and T cell reconstitution as compared to control H2B-GFP^neg^ HSCs (Figure 4C-D). Since H2B-GFP^neg^ HSCs were very rare in *Slc23a2*-deficient mice (Figure 3I), few of these cells could be isolated for transplantation. Nonetheless, both recipients that were transplanted with *Slc23a2*-deficient H2B-GFP^neg^ HSCs were long-term multilineage reconstituted by donor cells while only 1 of 9 recipients of control H2B-GFP^neg^ HSCs were long-term multilineage reconstituted by donor cells (Figure 4D). Ascorbate depletion thus increased the reconstituting potential of both frequently dividing and rarely dividing HSCs.

**Figure 4:**
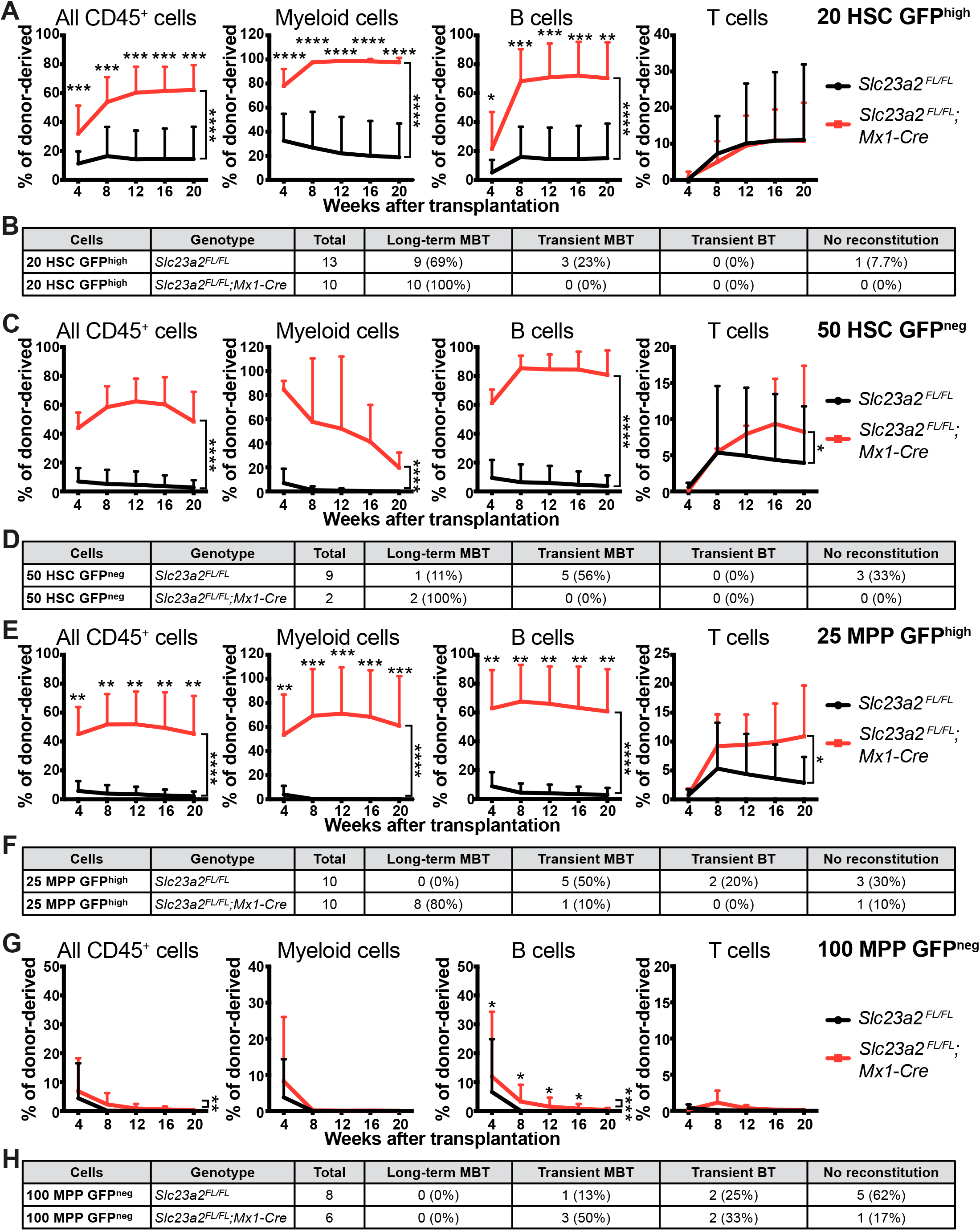
Ascorbate depletion confers long-term reconstituting potential upon quiescent MPPs. (A) Donor cell reconstitution in the blood of irradiated mice competitively transplanted with 20 H2B-GFP^high^ HSCs isolated from *Slc23a2*-deficient or littermate control mice (a total of 10-13 recipient mice per genotype transplanted with cells from 3 donors per genotype in 3 independent experiments). (B) Summary of the donor cell reconstitution profiles in panel A. (C) Donor cell reconstitution in mice competitively transplanted with 50 H2B-GFP^neg^ HSCs isolated from *Slc23a2*-deficient or littermate control mice (a total of 2-10 recipient mice per genotype transplanted with cells from 2 donors per genotype in 2 independent experiments). (D) Summary of the donor cell reconstitution profiles in panel C. (E) Donor cell reconstitution in mice competitively transplanted with 25 H2B-GFP^high^ MPPs isolated from *Slc23a2*-deficient or littermate control mice (a total of 10 recipient mice per genotype transplanted with cells from 2 donors per genotype in 2 independent experiments). (F) Summary of the donor cell reconstitution profiles in panel E. (G) Donor cell reconstitution in mice competitively transplanted with 100 H2B-GFP^neg^ MPPs isolated from *Slc23a2*-deficient or littermate control mice (a total of 6-8 recipient mice per genotype transplanted with cells from 2 donors per genotype in 2 independent experiments). (H) Summary of the donor cell reconstitution profiles in panel G. All data represent mean ± standard deviation. Statistical significance was assessed using Mann-Whitney tests to assess differences among genotypes at each timepoint and nparLD models with Holm-Sidak’s multiple comparisons corrections to test differences among genotypes in overall reconstitution (A, C, E, G). All statistical tests were two-sided (* *P*<0.05; ** *P*<0.01; *** *P*<0.001; **** *P*<0.0001).

We also competitively transplanted 25 H2B-GFP^high^ MPPs or 100 H2B-GFP^neg^ MPPs from *Slc23a2*-deficient and control mice into irradiated recipients. *Slc23a2*-deficient H2B-GFP^high^ MPPs gave significantly higher levels of donor myeloid, B, and T cell reconstitution than control H2B-GFP^high^ MPPs (Figure 4E): 8 of 10 recipients of *Slc23a2*-deficient H2B-GFP^high^ MPPs were long-term multilineage reconstituted by donor cells while none of 10 recipients of control H2B-GFP^high^ MPPs were long-term multilineage reconstituted (Figure 4F). *Slc23a2*-deficient H2B-GFP^neg^ MPPs gave significantly higher levels of donor B cell reconstitution than control H2B-GFP^neg^ MPPs (Figure 4G); however, none of the recipients of *Slc23a2*-deficient or control H2B-GFP^neg^ MPPs were long-term multilineage reconstituted (Figure 4H). Ascorbate depletion thus conferred long-term reconstituting potential upon quiescent, but not frequently dividing, MPPs.

### Ascorbate depletion does not promote MPP self-renewal via reduced Tet2 function

Ascorbate limits HSC reconstituting potential largely by promoting the function of Tet dioxygenases, particularly Tet2^18,40^. We thus tested whether ascorbate limited the self-renewal of MPPs by promoting Tet2 function. To do this, we generated *Slc23a2^FL/FL^;Tet2^FL/FL^;Mx1-Cre* double mutant mice, *Slc23a2^FL/FL^;Mx1-Cre* and *Tet2^FL/FL^;Mx1-Cre* single mutant mice, and *Slc23a2^FL/FL^* controls. *Tet2* and/or *Slc23a2* mutant mice did not significantly differ from control mice in terms of bone marrow cellularity, though bone marrow cellularity was significantly higher in *Tet2^FL/FL^;Mx1-Cre* as compared to *Slc23a2^FL/FL^;Mx1-Cre* mice (supplemental Figure 3A). *Slc23a2* deficiency significantly increased the frequency of MPP-1 cells but *Tet2* deficiency did not phenocopy the effect of *Slc23a2* deficiency (supplemental Figure 3B). This suggests that the effect of *Slc23a2* deficiency on MPP-1 frequency was not mediated by reduced *Tet2* function after ascorbate depletion.

We also competitively transplanted 50 MPP cells into irradiated recipient mice and assessed donor cell reconstitution. Only 1 of 8 recipients of *Slc23a2^FL/FL^* control MPPs were long-term multilineage reconstituted by donor cells; the remaining recipients were transiently multilineage reconstituted (supplemental Figure 3C-D). In contrast, 3 of 10 recipients of *Slc23a2* deficient MPPs were long-term multilineage reconstituted by donor cells (the remaining recipients were transiently multilineage reconstituted). *Tet2* deficiency did not phenocopy this increase in the frequency of long-term multilineage reconstituting MPPs as none of 11 recipients of *Tet2* deficient MPPs were long-term multilineage reconstituted by donor cells (10 of 11 recipients were transiently multilineage reconstituted). Thus, these data provided no evidence that the effects of *Slc23a2* deficiency on MPP reconstituting potential were mediated by reduced *Tet2* function. Consistent with this conclusion, *Slc23a2* and *Tet2* double knockout MPPs gave long-term multilineage reconstitution in 3 of 10 recipients, just as observed with *Slc23a2* single knockout MPPs. Thus, we found no evidence that *Tet2* deficiency accentuated the effect of *Slc23a2* deficiency on the reconstituting potential of MPPs.

## DISCUSSION

We reported previously that ascorbate limits HSC function during the reconstitution of irradiated mice^18^; however, the effect of ascorbate on MPP function had not been tested. In the current study, we characterized the effect of ascorbate depletion from hematopoietic cells by deleting the *Slc23a2* ascorbate transporter (Figure 1A). We were surprised to observe that this increased HSC and MPP function. Deletion of *Slc23a2* from hematopoietic cells increased quiescence (Figure 3E-I) reconstituting potential (Figure 2A-C and 2E-G) and self-renewal potential (Figure 2D and 2H-J) in both HSCs and MPPs. Ascorbate depletion was sufficient to confer MPPs with long-term self-renewal and reconstituting potential (Figure 2E-J).

Ascorbate depletion increased the reconstituting potential of both dividing and non-dividing HSCs (Figure 4A-D) but it only significantly increased the reconstituting potential of the non-dividing (or slowly dividing) fraction of MPPs (Figure 4E-H). When combined with the recent observation of higher ascorbate levels in dividing as compared to quiescent HSCs^21^, these observations suggest that ascorbate promotes cell division by HSCs and MPPs, at least under steady-state conditions. Nonetheless, when compared to control cells, ascorbate depleted HSCs and MPPs gave higher levels of donor cell reconstitution upon transplantation into irradiated mice; therefore, ascorbate depleted HSCs and MPPs are capable of repeated cell divisions in myeloablated mice even if they are more quiescent under steady-state conditions.

Most of the effect of ascorbate on HSC function is mediated by Tet2^18^. Superphysiological ascorbate levels attenuate the effects of *Tet2* deficiency in mouse and human HSCs^40,41^, partly by promoting Tet3 function^40^. Furthermore, ascorbate promotes plasma cell differentiation by increasing Tet2 and Tet3 function^28^. However, the effects of ascorbate depletion on MPP reconstituting potential do not appear to be mediated by Tet2 because *Tet2* deficiency did not phenocopy or exacerbate the effects of ascorbate depletion on MPPs (supplemental Figure 3A-D). Many cell cycle-related genes were reduced in expression in ascorbate depleted as compared to control MPPs (Figure 3C and 3D), consistent with the increased quiescence of *Slc23a2*-deficient MPPs. However, the mechanism by which ascorbate promotes cell division and limits self-renewal potential in MPPs remains unclear.

## ACKNOWLEDGMENTS

S.J.M. is a Howard Hughes Medical Institute Investigator, the Mary McDermott Cook Chair in Pediatric Genetics, the Kathryn and Gene Bishop Distinguished Chair in Pediatric Research, the director of the Hamon Laboratory for Stem Cells and Cancer, and a Cancer Prevention and Research Institute of Texas Scholar. This work was supported by the National Institutes of Health (DK118745) and the Kleberg Foundation. S.C. was supported by an EMBO Long-Term Fellowship (ALTF 722-2015), E.C.J. was supported by a postdoctoral fellowship from the Damon Runyon Cancer Research Foundation (2278-16), and A.W.D. was supported by a Ruth L. Kirschstein National Research Service Award Postdoctoral fellowship from the National Heart Lung and Blood Institute (F32HL135975). We thank Jian Xu and Michalis Agathocleous for helpful discussions. We thank M. Ortiz and the Moody Foundation Flow Cytometry Facility, T. Sharma and the Mouse Genome Engineering Facility, and the Metabolomics Facility in Children’s Medical Center Research Institute at UT Southwestern. We also thank M. Mulkey for mouse colony management and the BioHPC High-Performance Facility Cloud at UT Southwestern Medical Center for providing computational resources.

## AUTHORSHIP CONTRIBUTIONS

S.C. and S.J.M. conceived the project, designed and interpreted experiments. S.C. performed most of the experiments, with technical assistance from D.C., A.W.D., E.C.J., B.D. and A.R. B.C. and Z.Z. performed the RNA-Seq and statistical analyses. S.C. and S.J.M. wrote the manuscript.

