## Supplemental data for "Ascorbate depletion increases quiescence and self-renewal potential in hematopoietic stem cells and multipotent progenitors"

### SUPPLEMENTAL MATERIAL

**Supplemental Table 1. Cell populations analyzed by flow cytometry in this study.**

| Population | Abbrev. | Markers | Reference |
|---|---|---|---|
| Hematopoietic Stem Cells | HSC | CD150$^{+}$ CD48$^{-}$ Lin$^{-}$ Sca1$^{+}$ c-kit$^{+}$ | [1] |
| Multipotent progenitors | MPP | CD150$^{-}$ CD48$^{-}$ Lin$^{-}$ Sca1$^{+}$ c-kit$^{+}$ | [1] |
| HPC1 progenitors | HPC1 | CD150$^{-}$ CD48$^{+}$ Lin$^{-}$ Sca1$^{+}$ c-kit$^{+}$ | [2] |
| HPC2 progenitors | HPC2 | CD150$^{+}$ CD48$^{+}$ Lin$^{-}$ Sca1$^{+}$ c-kit$^{+}$ | [2] |
| HSC-1 subpopulation | HSC-1 | CD150$^{+}$ CD48$^{-}$ Lin$^{-}$ Sca1$^{+}$ c-kit$^{+}$ CD229$^{-}$ CD244$^{-}$ | [2] |
| HSC-2 subpopulation | HSC-2 | CD150$^{+}$ CD48$^{-}$ Lin$^{-}$ Sca1$^{+}$ c-kit$^{+}$ CD229$^{-}$ CD244$^{+}$ | [2] |
| MPP-1 subpopulation | MPP-1 | CD150$^{-}$ CD48$^{-}$ Lin$^{-}$ Sca1$^{+}$ c-kit$^{+}$ CD229$^{-}$ CD244$^{-}$ | [2] |
| MPP-2 subpopulation | MPP-2 | CD150$^{-}$ CD48$^{-}$ Lin$^{-}$ Sca1$^{+}$ c-kit$^{+}$ CD229$^{-}$ CD244$^{+}$ | [2] |
| MPP-3 subpopulation | MPP-3 | CD150$^{-}$ CD48$^{-}$ Lin$^{-}$ Sca1$^{+}$ c-kit$^{+}$ CD229$^{+}$ CD244$^{+}$ | [2] |
| Common Lymphoid Progenitors | CLP | Lin$^{-}$ c-kit$^{low}$ Sca1$^{low}$ Flt3$^{+}$ IL7R$\alpha^{+}$ | [3] |
| Common Myeloid Progenitors | CMP | Lin$^{-}$ c-kit$^{+}$ Sca1$^{-}$ CD34$^{+}$ CD16/32$^{-}$ | [4] |
| Megakaryocyte-erythrocyte progenitors | MEP | Lin$^{-}$ c-kit$^{+}$ Sca1$^{-}$ CD34$^{-}$ CD16/32$^{-}$ | [4] |
| Granulocyte-monocyte progenitors | GMP | Lin$^{-}$ c-kit$^{+}$ Sca1$^{-}$ CD34$^{+}$ CD16/32$^{+\ -}$ | [4] |
| Pre-Megakaryocyte-erythrocyte progenitors | PreMegE | Lin$^{-}$ c-kit$^{+}$ Sca1$^{-}$ CD41$^{-}$ CD16/32$^{-}$ CD150$^{+}$ CD105$^{-}$ | [5] |
| PreCFU-E progenitors | PreCFU-E | Lin$^{-}$ c-kit$^{+}$ Sca1$^{-}$ CD41$^{-}$ CD16/32$^{-}$ CD150$^{+}$ CD105$^{+}$ | [5] |
| CFU-E progenitors | CFU-E | Lin$^{-}$ c-kit$^{+}$ Sca1$^{-}$ CD41$^{-}$ CD16/32$^{-}$ CD150$^{-}$ CD105$^{+}$ | [5] |
| B cells | | B220$^{+}$ CD3$^{-}$ Mac1$^{-}$ | |
| T cells | | CD3$^{+}$ B220$^{-}$ Mac1$^{-}$ | |
| Granulocytes | | Mac1$^{+}$Gr1$^{+}$ B220$^{-}$ CD3$^{-}$ | |

**Supplemental Table 2. Ten most upregulated and downregulated genes in *Slc23a2* deficient as compared to control HSCs and MPPs**

A: Ten most upregulated genes in *Slc23a2*-deficient as compared to control HSCs

| Gene symbol | Gene name | Fold change | FDR |
|---|---|---|---|
| Paqr5 | progestin and adipoQ receptor family member V | 2.64 | 1.76E-05 |
| H19 | H19, imprinted maternally expressed transcript | 2.58 | 0.0028 |
| Adamtsl5 | ADAMTS-like 5 | 2.37 | 5.89E-05 |
| Efnb2 | ephrin B2 | 2.32 | 0.00038 |
| Ltbp4 | latent transforming growth factor beta binding protein 4 | 2.19 | 0.0025 |
| Hspa2 | heat shock protein 2 | 2.06 | 2.77E-05 |
| Pim2 | proviral integration site 2 | 2.05 | 2.48E-14 |
| Slc7a7 | solute carrier family 7 (cationic amino acid transporter, y+ system), member 7 | 2.03 | 0.0043 |
| Setbp1 | SET binding protein 1 | 2.01 | 2.70E-05 |

B: Ten most downregulated genes in *Slc23a2*-deficient as compared to control HSCs

| Gene symbol | Gene name | Fold change | FDR |
|---|---|---|---|
| Gpx7 | glutathione peroxidase 7 | 0.19 | 1.34E-08 |
| Gas6 | growth arrest specific 6 | 0.26 | 8.89E-13 |
| Ly9 | lymphocyte antigen 9 | 0.28 | 3.38E-09 |
| Gstm7 | glutathione S-transferase, mu 7 | 0.28 | 2.23E-06 |
| Arap3 | ArfGAP with RhoGAP domain, ankyrin repeat and PH domain 3 | 0.28 | 2.84E-10 |
| Fbp1 | fructose bisphosphatase 1 | 0.29 | 1.59E-05 |
| Depdc1a | DEP domain containing 1a | 0.34 | 1.82E-07 |
| Slc23a2 | solute carrier family 23 (nucleobase transporters), member 2 | 0.37 | 1.09E-26 |
| Kif14 | kinesin family member 14 | 0.37 | 1.04E-07 |
| Pimreg | PICALM interacting mitotic regulator | 0.38 | 1.23E-05 |

C: Ten most upregulated genes in *Slc23a2*-deficient as compared to control MPPs

| Gene symbol | Gene name | Fold change | FDR |
|---|---|---|---|
| Rag1 | recombination activating gene 1 | 4.88 | 3.23E-07 |
| Igll1 | immunoglobulin lambda-like polypeptide 1 | 4.55 | 1.99E-13 |

| | | | |
|---|---|---|---|
| Dok3 | docking protein 3 | 4.47 | 3.40E-09 |
| Vpreb1 | pre-B lymphocyte gene 1 | 3.82 | 0.00067 |
| Scube3 | signal peptide, CUB domain, EGF-like 3 | 3.20 | 1.25E-07 |
| Mpo | myeloperoxidase | 3.05 | 4.48E-06 |
| Tjp1 | tight junction protein 1 | 2.77 | 9.75E-06 |
| Gm3448 | predicted gene 3448 | 2.76 | 0.0013 |
| Gdpd3 | glycerophosphodiester phosphodiesterase domain containing 3 | 2.73 | 2.63E-05 |
| Akap12 | A kinase (PRKA) anchor protein (gravin) 12 | 2.62 | 0.0047 |

D: Ten most downregulated genes in *Slc23a2*-deficient as compared to control MPPs

| **Gene symbol** | **Gene name** | **Fold change** | **FDR** |
|---|---|---|---|
| Fbp1 | fructose bisphosphatase 1 | 0.18 | 3.01E-09 |
| Tgfbi | transforming growth factor, beta induced | 0.20 | 4.55E-20 |
| Gstm7 | glutathione S-transferase, mu 7 | 0.26 | 0.0018 |
| Kif18b | kinesin family member 18B | 0.27 | 5.33E-05 |
| Nuf2 | NUF2, NDC80 kinetochore complex component | 0.27 | 7.45E-07 |
| Kntc1 | kinetochore associated 1 | 0.27 | 1.47E-09 |
| Gpx7 | glutathione peroxidase 7 | 0.28 | 0.00141271 |
| Arap3 | ArfGAP with RhoGAP domain, ankyrin repeat and PH domain 3 | 0.29 | 1.01E-09 |
| Card11 | caspase recruitment domain family, member 11 | 0.30 | 7.94E-06 |
| Bard1 | BRCA1 associated RING domain 1 | 0.30 | 1.79E-05 |

### SUPPLEMENTAL FIGURE LEGENDS

**Supplemental Figure 1. Targeting strategy to generate the *Slc23a2*$^{FL}$ allele and representative flow cytometry gates used to identify hematopoietic stem and progenitor cell populations.** (A) The donor ssDNA contained *loxP* sequences on both sides of exon 5. The *loxP* insertion sites were chosen to avoid disrupting sequences conserved among species and to cause a frameshift upon Cre-mediated recombination. CRISPR/Cas9-mediated recombination was performed by injecting donor ssDNA, Cas9 protein, tracrRNA and 2 sgRNAs targeting the endogenous sequences where *loxP* sites were inserted into C57BL/6 zygotes. (B) Correctly targeted founder mice were identified by long-range PCR with primers flanking the whole region targeted by donor ssDNA followed by Sanger sequencing. (C) PCR genotyping. (D-F) Gating strategy to isolate HSCs, MPPs, HPC1 cells, HPC2 cells, MEPs, CMPs, GMPs (D) PreMegEs, PreCFU-Es and CFU-Es (E), and CLPs (F) from bone marrow by flow cytometry[1,3-5] (see also supplemental Table 1 for a list of the markers used to identify each cell population).

**Supplemental Figure 2. Ascorbate depletion did not confer long-term reconstituting potential upon HPC1 or HPC2 cells.** (A) Donor cell reconstitution in the blood of irradiated mice competitively transplanted with 500 HPC1 cells isolated from *Slc23a2*-deficient or littermate control mice (a total of 10-13 recipient mice per genotype transplanted with cells from 3 donors per genotype in 3 independent experiments). (B) Summary of the donor cell reconstitution profiles in panel A. (C) Donor cell reconstitution in the blood of irradiated mice competitively transplanted with 200 HPC2 cells isolated from *Slc23a2*-deficient or littermate control mice (a total of 7 recipient mice per genotype transplanted with cells from 2 donors per genotype in 2 independent experiments). (D) Summary of the donor cell reconstitution profiles in panel C. All data represent mean ± standard deviation. Statistical significance was assessed using Mann-Whitney tests to assess differences among genotypes at each timepoint and nparLD models with Holm-Sidak’s multiple comparisons corrections to assess differences among genotypes in overall reconstitution (A and C). All statistical tests were two-sided (* *P*<0.05; ** *P*<0.01; *** *P*<0.001).

**Supplemental Figure 3. Tet2 does not appear to mediate the effects of ascorbate depletion on the reconstituting potential of MPPs.** Each panel in this figure compares cells from control mice, *Slc23a2*$^{FL/FL}$*;Mx1-Cre* mice, *Tet2*$^{FL/FL}$*;Mx1-Cre* mice, and *Slc23a2*$^{FL/FL}$*;Tet2*$^{FL/FL}$*;Mx1-Cre* mice. (A) Bone marrow cellularity in one tibia and one femur. (B) Frequencies of MPP-1, MPP-2, and MPP-3 cells in the bone marrow (panels A and B show a total of 4-5 mice per genotype analyzed in 4 independent experiments). (C) Donor cell reconstitution in the blood of irradiated mice competitively transplanted with 50 MPPs (a total of 8-11 recipient mice per genotype transplanted with cells from 3 donors per genotype in 3 independent experiments). (D) Summary of the donor cell reconstitution profiles in panel C. All data represent mean ± standard deviation. Each dot represents a different mouse in panels A and B. Statistical significance was assessed using one-way ANOVAs with Tukey's correction for multiple comparisons (A-B), Kruskal-Wallis tests to assess differences among genotypes within each timepoint and nparLD models with Holm-Sidak's multiple comparisons corrections to test differences among genotypes in overall reconstitution (C). All statistical tests were two-sided (* *P*<0.05; ** *P*<0.01; *** *P*<0.001; **** *P*<0.0001).

Comazzetto et al. Supplemental Figure 1

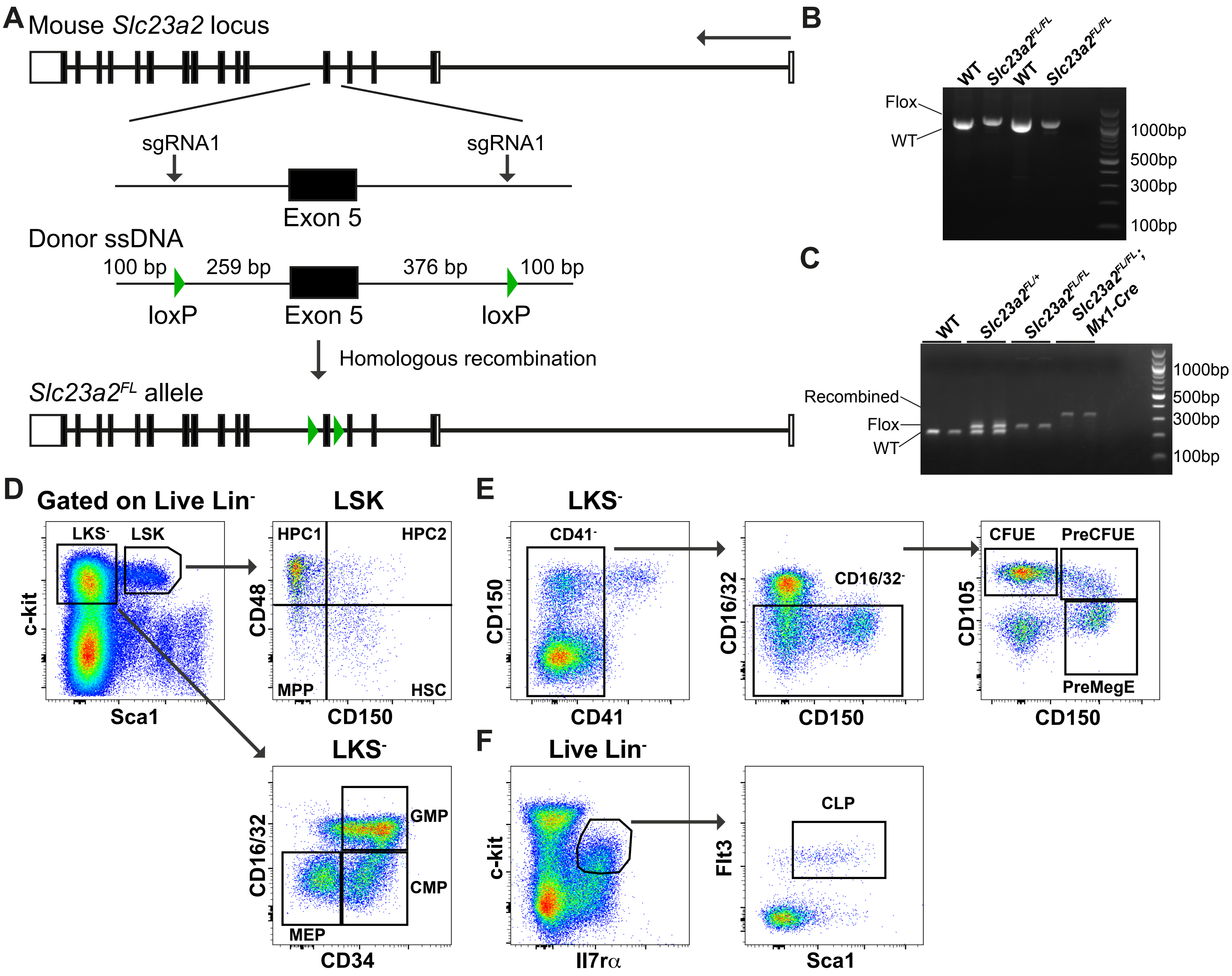

Comazzetto et al. Supplemental Figure 2

**A**

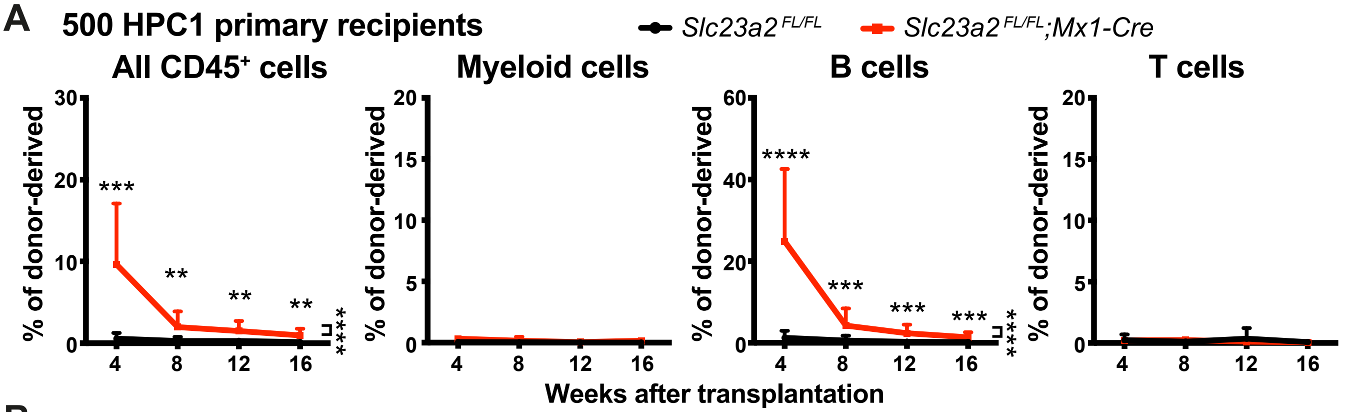


**B**

| Cells | Genotype | Total | Long-term MBT | Transient MB | Transient BT | No recon-stitution |
|---|---|---|---|---|---|---|
| **500 HPC1** | *Slc23a2^FL/FL^* | 13 | 0 (0%) | 0 (%) | 7 (54%) | 6 (46%) |
| **500 HPC1** | *Slc23a2^FL/FL^;Mx1-Cre* | 10 | 0 (0%) | 1 (10%) | 9 (90%) | 0 (0%) |

**C**

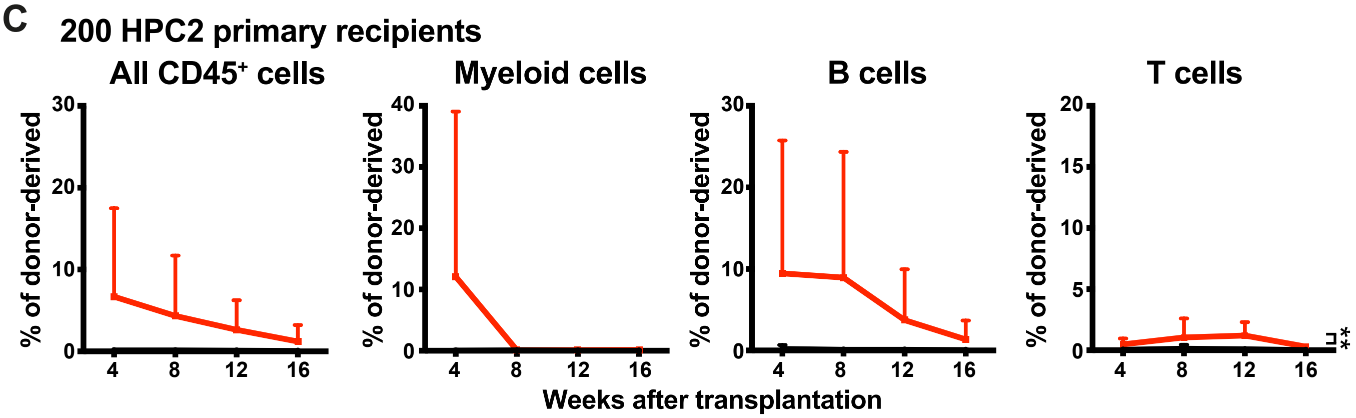


**D**

| Cells | Genotype | Total | Long-term MBT | Transient MBT | Transient BT | No recon-stitution |
|---|---|---|---|---|---|---|
| **200 HPC2** | *Slc23a2^FL/FL^* | 7 | 0 (0%) | 0 (0%) | 2 (29%) | 5 (71%) |
| **200 HPC2** | *Slc23a2^FL/FL^;Mx1-Cre* | 7 | 0 (0%) | 2 (29%) | 2 (29%) | 3 (42%) |

Comazzetto et al. Supplemental Figure 3

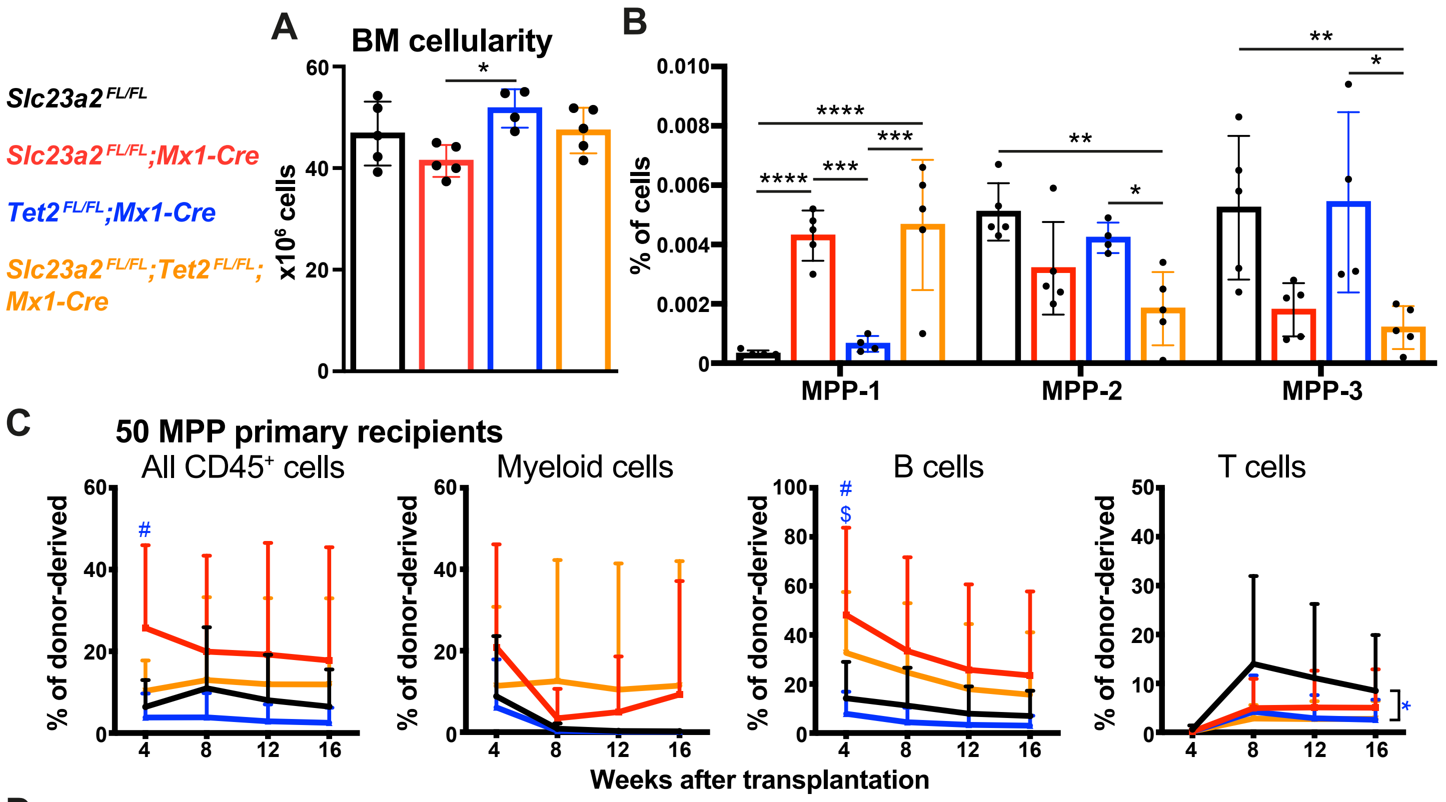


**D**

| Cells | Genotype | Total | Long-term MBT | Transient MBT | Transient BT | No recon-stitution |
|---|---|---|---|---|---|---|
| **50 MPP** | *Slc23a2$^{FL/FL}$* | 8 | 1 (13%) | 4 (50%) | 3 (37%) | 0 (0%) |
| **50 MPP** | *Slc23a2$^{FL/FL}$;Mx1-Cre* | 10 | 3 (30%) | 4 (40%) | 3 (30%) | 0 (0%) |
| **50 MPP** | *Tet2$^{FL/FL}$;Mx1-Cre* | 11 | 0 (0%) | 6 (55%) | 4 (36%) | 1 (9.1%) |
| **50 MPP** | *Slc23a2$^{FL/FL}$;Tet2$^{FL/FL;}$Mx1-Cre* | 10 | 3 (30%) | 4 (40%) | 3 (30%) | 0 (0%) |
